## Supporting Information for "Genetic drift promotes and recombination hinders speciation on holey fitness landscapes"

**S1 Fig. Genetic drift promotes the accumulation of genetic incompatibilities in the RNA folding model without intrinsic fitness differences between genotypes.**

**S2 Fig. Recombination hinders the accumulation of genetic incompatibilities in the RNA folding model without intrinsic fitness differences between genotypes.**

**S3 Fig. Number of divergent alleles from one population that have not have been tested by natural selection in the genetic background of another population.**

**S1 Text. Number of divergent alleles from one population that have not have been tested by natural selection in the genetic background of another population.**

**S1 Table. Synonymous nucleotide diversity ($\pi_s$) at the Adh locus for different species of *Drosophila*.**

**S2 Table. Total recombination map length for different species of *Drosophila*.**

**S3 Table. Phylogenetic signal for each trait analyzed in Fig. 10.**

**S4 Table. Asymmetry in synonymous nucleotide diversity ($\pi_s$) at the Adh locus and in postzygotic reproductive isolation (RI) for different species of *Drosophila*.**

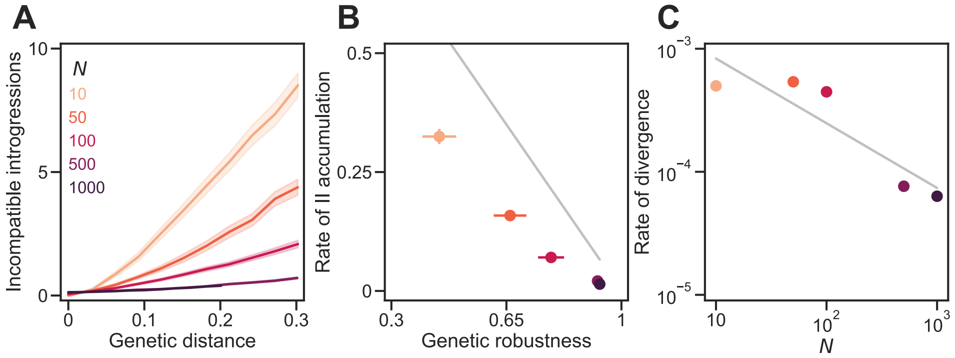


**Fig S1. Genetic drift promotes the accumulation of genetic incompatibilities in the RNA folding model without intrinsic fitness differences between viable genotypes. (A)** IIs accumulated approximately linearly with $D$ in sexual populations of different sizes ($N$). In all simulations, populations evolved under the RNA folding model (with $\sigma = 0$ and $L = 100$) and experienced $U = 0.1$, random mating, and free recombination. **(B)** IIs accumulated faster in smaller populations because they evolved lower $\nu$. The gray line shows $b/L = 1 - \nu$ (see Eq 1). **(C)** Large populations diverged more slowly. The gray line shows a power law with exponent $-0.5$. Plotted values in all panels are means of 200 replicate simulation runs at each $N$. Error bands in (A) and bars in (B) are 95% CIs (in (C) they are hidden by the points). See Fig 7 for more details.

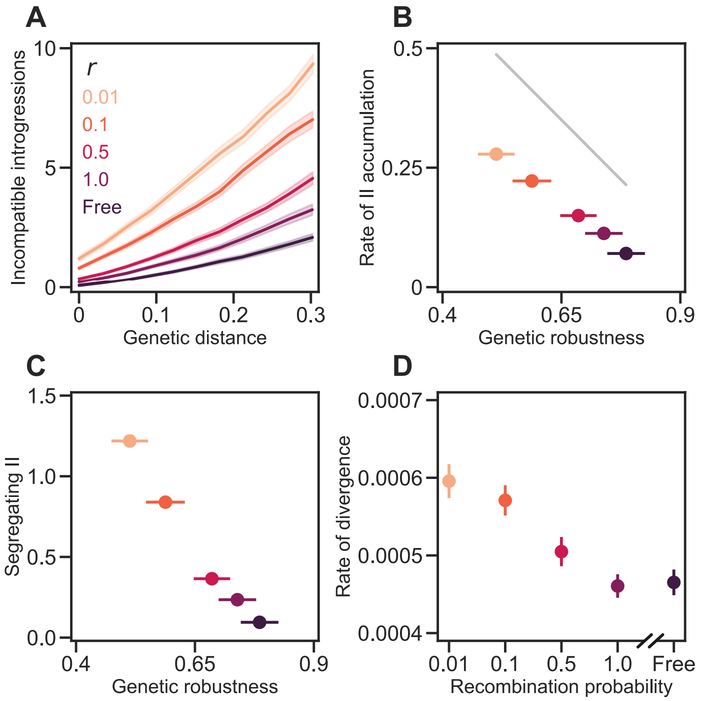


**Fig S2. Recombination hinders the accumulation of genetic incompatibilities in the RNA folding model without intrinsic fitness differences between viable genotypes.** **(A)** IIs accumulated approximately linearly with $D$ in populations of $N = 100$ individuals experiencing different recombination probabilities. In all simulations, populations evolved under the RNA folding model (with $\sigma = 0$ and $L = 100$) and experienced $U = 0.1$ and random mating. **(B)** IIs accumulated faster in populations experiencing low recombination probability because they evolved lower $\nu$. The gray line shows $b/L = 1 - \nu$ (see Eq 1). **(C)** Populations experiencing low recombination probability accumulated more segregating IIs. **(D)** Populations experiencing high $r$ diverged more slowly. Values show the rate of increase in $D$ per generation. Plotted values in all panels are means of 200 replicate simulation runs at each $r$. Error bands in (A) and bars in (B)–(D) are 95% CIs (some are hidden by the points). See Fig 8 for more details.

### S1 Text. Number of divergent alleles from one population that have not have been tested by natural selection in the genetic background of another population

If two populations evolving in allopatry have diverged at $k$ diallelic loci and there have been no reversals, then $k-1$ divergent alleles from one population will not have been tested by natural selection in the genetic background of the other population. The number of alleles is $k-1$ rather than $k$ because we must exclude one allele that *has* been tested in the recipient background. This point explains both the $k-1$ and $\tilde{\nu}$ terms in Eq 1. Which allele is to be excluded depends on the number of substitutions that have taken place in the recipient population. There are two possible scenarios, A and B (S3 Fig).

**Scenario A:** If no substitutions have taken place in the recipient population we exclude the introgression of the derived allele at the first locus to undergo a substitution in the donor population because the resulting genotype has been tested by natural selection (S3A Fig, dashed arrow).

**Scenario B:** If at least one substitution has taken place in the recipient population we exclude the introgression of the ancestral allele at the last locus to undergo a substitution in the recipient population because the resulting genotype has been tested by natural selection (S3B Fig, dashed arrow; also, Fig 1B).

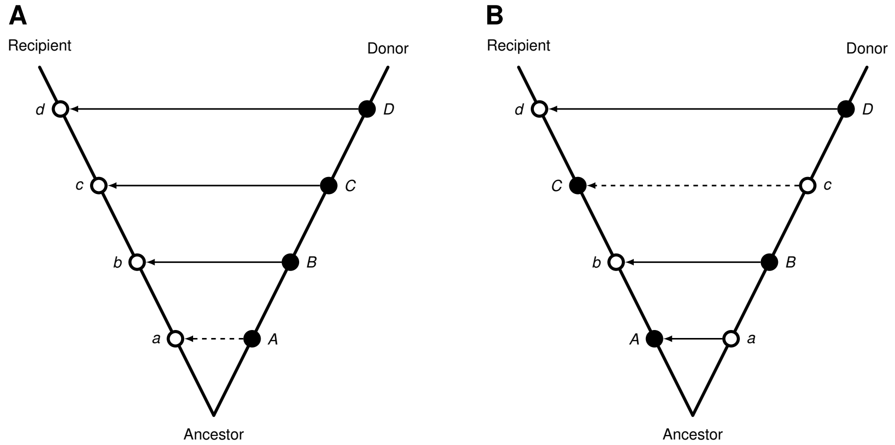


**Fig S3. Number of divergent alleles from one population that have not have been tested by natural selection in the genetic background of another population. (A)** Example of scenario A. Two populations diverge in allopatry. Both populations are initially fixed for lowercase alleles (open circles) at four loci *(abcd)*. Derived alleles are indicated by uppercase letters (closed circles). The donor population undergoes four substitutions, fixing the *ABCD* genotype. The recipient population does not undergo any substitutions, retaining the ancestral genotype *abcd*. Arrows indicate introgressions of divergent alleles from the donor population to the recipient population. Three introgressed genotypes have not been tested by natural selection (*aBcd*, *abCd*, and *abcD*; solid arrows) but one has (*Abcd*, dashed arrow). **(B)** Example of scenario B. Each population undergoes two substitutions, the recipient population fixing the *AbCd* genotype and the donor population fixing the *aBcD* genotype. Three introgressed genotypes have not been tested by natural selection (*abCd*, *ABCd*, and *AbCD*; solid arrows) but one has (*Abcd*, dashed arrow).

**S1 Table.** Synonymous nucleotide diversity (πS) at the *Adh* locus for different species of *Drosophila* .

| group | species | πS | sequences | reference | notes |
|---|---|---|---|---|---|
| *ananassae* | *ananassae* | 0.015822 | 10 | 1 | |
| *obscura* | *bogotana* | 0.004841 | 8 | 2 | |
| *obscura* | *miranda* | 0.003703 | 12 | 3 | |
| *obscura* | *persimilis* | 0.013317 | 6 | 4 | |
| *obscura* | *pseudoobscura* | 0.016660 | 139 | 2, 5, 6 | |
| *obscura* | *subobscura* | 0.023465 | 16 | 7 | |
| *virilis* | *americana* | 0.034155 | 19 | 8 | G96X |
| *virilis* | *americana* | 0.023301 | 19 | 8 | G96Y |
| *virilis* | *borealis* | 0.059733 | 2 | 9, 10 | |
| *virilis* | *lummei* | 0.000000 | 2 | 9, 11 | |
| *virilis* | *texana* | 0.029706 | 10 | 8 | |
| *virilis* | *virilis* | 0.005490 | 2 | 11 | |
| *repleta* | *arizonae* | 0.013155 | 13 | 12 | Adh-1 |
| *repleta* | *arizonae* | 0.052620 | 11 | 12 | Adh-2 |
| *repleta* | *buzzatii* | 0.016323 | 4 | 13 | |
| *repleta* | *mojavensis* | 0.013141 | 13 | 12 | Adh-1 |
| *repleta* | *mojavensis* | 0.012597 | 15 | 14 | Adh-1 |
| *repleta* | *mojavensis* | 0.021064 | 13 | 12 | Adh-2 |
| *repleta* | *mojavensis* | 0.009859 | 15 | 14 | Adh-2 |
| *repleta* | *hydei* | 0.017564 | 9 | 13 | |
| *repleta* | *mulleri* | 0.019467 | 6 | 13 | |
| *immigrans* | *albomicans* | 0.031127 | 16 | 15 | |
| *immigrans* | *hypocausta* | 0.031060 | 2 | 16 | |
| *immigrans* | *nasuta* | 0.003687 | 16 | 15 | |
| *immigrans* | *siamana* | 0.038144 | 2 | 16, 17 | |
| *melanogaster* | *lutescens* | 0.009242 | 4 | 18–21 | |
| *melanogaster* | *mauritiana* | 0.002228 | 6 | 22 | |
| *melanogaster* | *melanogaster* | 0.024036 | 10 | 23 | |
| *melanogaster* | *melanogaster* | 0.027122 | 4 | 24 | |
| *melanogaster* | *melanogaster* | 0.030007 | 11 | 25 | |
| *melanogaster* | *sechellia* | 0.000000 | 2 | 22 | |
| *melanogaster* | *simulans* | 0.027407 | 7 | 26 | |
| *melanogaster* | *yakuba* | 0.007072 | 12 | 26 | |
| *melanogaster* | *yakuba* | 0.015629 | 36 | 27 | |
| *willistoni* | *equinoxialis* | 0.005490 | 2 | 28 | |
| *willistoni* | *nebulosa* | 0.011506 | 4 | 28, 29 | |
| *willistoni* | *paulistorum* | 0.018475 | 3 | 28 | |
| *willistoni* | *willistoni* | 0.018035 | 18 | 30, 31 | |
| *montium* | *auraria* | 0.045728 | 2 | 32 | |

| | | | | |
|---|---|---|---|---|
| *montium* | *birchii* | 0.051745 | 2 | 32 |
| *montium* | *kikkawai* | 0.034314 | 21 | 33 |
| *montium* | *leontia* | 0.027858 | 2 | 32 |
| *montium* | *lini* | 0.062418 | 2 | 32 |
| *montium* | *rufa* | 0.000000 | 2 | 32 |
| *montium* | *serrata* | 0.022432 | 2 | 32 |
| *montium* | *triauraria* | 0.011194 | 2 | 32 |
| *montium* | *triauraria* | 0.019847 | 3 | 34 |

**Methods:** Sequences were obtained from GenBank. Synonymous nucleotide diversity (πS) was calculated using the Nei-Gojobori method in MEGA-CC version 11.0.13 (35).

**References:** 1) Shih & Jones 2008 Genetics 180: 1261–1263. 2) Shaeffer & Miller 1991 PNAS 88: 6097–6101. 3) Yi et al. 2003 Genetics 164: 1369–1381. 4) Wang et al. 1997 Genetics 147: 1091–1106. 5) Schaeffer & Miller 1993 Genetics 135: 541–552. 6) Schaeffer 2002 Genet. Res. 80: 163–175. 7) Jones et al. 2005 Genetics 170: 207–219. 8) McAllister & Charlesworth 1999 Genetics 153: 221–233. 9) Nurminsky et al. 1996 Mol. Biol. Evol. 13: 132–149. 10) Morales-Hojas 2011 Mol. Phyl. Evol. 60: 249–258. 11) Wang et al. 2006 Mol. Phyl. Evol. 40: 484–500. 12) Matzkin & Eanes 2003 Genetics 163: 181–194. 13) Begun 1997 Genetics 145: 375–382. 14) Matzkin 2004 Mol. Biol. Evol. 21: 276–285. 15) Satomura & Tamura 2016 Mol. Biol. Evol. 33: 367–374. 16) Katoh et al. 2007 Zool. Sci. 24: 913–921. 17) Rice et al. 2018 Evol. Dev. 20: 78–88. 18) Katoh et al. 2000 J. Mol. Evol. 51: 122–130. 19) O'Grady & Kidwell 2002 Mol. Phyl. Evol. 22: 442–453. 20) Ko, David & Akashi 2003 J. Mol. Evol. 57:  562–573. 21) Katoh & Watada 2015 GenBank: LC057203.1. 22) Kliman et al. 2000 Genetics 156: 1913–1931. 23) Begun et al. 1999 Mol. Biol. Evol. 16: 1816–1819. 24) Laurie et al. 1991 Genetics 129: 489–499. 25) Kreitman 1983 Nature 304: 412–417. 26) McDonald & Kreitman 1991 Nature 351: 652–654. 27) Siddiq et al. 2017 Nat. Ecol. Evol. 1: 0025. 28) Gleason et al. 1998 Evolution 52: 1093–1103. 29) Gao et al. 2011 Mol. Phyl. Evol. 60: 98–107. 30) Anderson et al. 1993 Mol. Biol. Evol. 10: 605–618. 31) Griffith & Powell 1997 J. Mol. Evol. 45: 232–237. 32) Chen et al. 2013 Zool. Sci. 30: 1056–1062. 33) Goto et al. 2004 Genes Genet. Syst. 19–26. 34) Dai, Lu, Lv, Chen, Cheng & Zhang 2001 GenBank: AF348879.1, AF348880.1, AF348882.1. 35) Kumar et al. 2012 Bioinformatics 28: 2685–2686.

**S2 Table.** Total recombination map length for different species of *Drosophila* .

| group | species | map length (cM) | reference | mapping function | corrected† |
|---|---|---|---|---|---|
| *ananassae* | *ananassae* | 350.1 | 1 | Kosambi | yes |
| *immigrans-tripunctata* | *funebris* | 895.6 | 1 | Kosambi | yes |
| *immigrans-tripunctata* | *mediopunctata* | 609.6 | 1 | Kosambi | yes |
| *melanogaster* | *mauritiana* | 488.5 | 2 | Foss | yes |
| *melanogaster* | *melanogaster* | 294.9 | 1 | Kosambi | yes |
| *melanogaster* | *melanogaster* | 287.3 | 3 | N/A* | no |
| *melanogaster* | *simulans* | 426.2 | 4 | Kosambi | yes |
| *montium* | *serrata* | 282.3 | 5 | Kosambi | yes |
| *obscura* | *persimilis* | 605.1 | 1 | Kosambi | yes |
| *obscura* | *pseudoobscura* | 557.1 | 1 | Kosambi | yes |
| *obscura* | *subobscura* | 1007.6 | 1 | Kosambi | yes |
| *repleta* | *buzzatii* | 696.5 | 1 | Kosambi | yes |
| *repleta* | *hydei* | 655.2 | 1 | Kosambi | yes |
| *virilis* | *montana* | 632.7 | 6 | Kosambi | yes |
| *virilis* | *virilis* | 732.3 | 7 | N/A* | no |
| *willistoni* | *willistoni* | 320.4 | 8 | Kosambi | yes |

* High resolution mapping.

† The length of each chromosome was multiplied by ($n$ + 1) / ($n$ – 1), where $n$ is the number of markers per chromosome.

**References:** 1) Cáceres et al. 1999 Genetics 153: 251–259. 2) True et al. 1996 Genetics 142: 507–523. 3) Comeron et al. 2012 PLoS Genetics 8: e1002905. 4) Barker & Moth 2001 Dros. Inf. Serv. 84: 205–206. 5) Stocker et al. 2012 G3 2: 287–297. 6) Schäfer et al. 2010 J. Evol. Biol. 23: 518–527. 7) Hemmer et al. 2020 Mobile DNA 11: 10. 8) Spassky & Dobzhansky 1950 Heredity 4: 201–215.

**S3 Table.** Phylogenetic signal for each trait analyzed in Fig. 10.

| trait | Blomberg's *K* | *P*-value | Pagel's *λ* | *P*(*λ*=0) | *P*(*λ*=1) |
|---|---|---|---|---|---|
| nucleotide diversity | 0.430 | 0.928 | 0.000 | 1.000 | 0.018 |
| map length | 1.124 | 0.056 | 1.155 | 0.156 | 0.175 |
| postzygotic RI velocity | 0.427 | 0.957 | 0.000 | 1.000 | 0.017 |

**Methods:** The phylogeny shown in Fig. 10A was used for all analyses.
Blomberg's *K* and corresponding *P*-value were calculated using the R package `phytools` version 1.0-3 (1). The *P*-value was calculated from 100,000 randomizations.
Pagel's λ was estimated by maximum likelihood using `phytools`. *P* values were calculated by likelihood ratio tests. The log-likelihood for *λ*=0 was obtainec from `phytools`; the log-likelihood for *λ*=1 was calculated using the R package `geiger` version 2.0.10 (2).

**References:** 1) Revell 2012 Meth. Ecol. Evol. 3: 217–223. 2) Pennell et al. 2014 Bioinformatics 30: 2216–2218.

**S4 Table.** Asymmetry in synonymous nucleotide diversity (πS) at the *Adh* locus and in postzygotic reproductive isolation (RI) for different species of *Drosophila* .

| species 1 | species 2 | group | πS1 | πS2 | πS asym | RI asym | RI12* | RI21* | references | notes |
|---|---|---|---|---|---|---|---|---|---|---|
| *arizonae* | *mojavensis* | *repleta* | 0.032887 | 0.014165 | 0.018722 | 0.334 | 0.500 | 0.166 | | mean of 4 estimates |
| *auraria* | *triauraria* | *montium* | 0.045728 | 0.015520 | 0.030208 | 1.000 | 1.000 | 0.000 | | |
| *auraria* | *subauraria* | *montium* | 0.045728 | 0.011091 | 0.034637 | 0.500 | 1.000 | 0.500 | 1 | |
| *equinoxialis* | *paulistorum* | *willistoni* | 0.005489 | 0.018475 | -0.012986 | -0.500 | 0.500 | 1.000 | | |
| *lini* | *ogumai* | *montium* | 0.062418 | 0.022183 | 0.040235 | 0.000 | 0.500 | 0.500 | 1 | |
| *lini* | *ohnishii* | *montium* | 0.062418 | 0.027833 | 0.034585 | 0.000 | 0.500 | 0.500 | 1 | |
| *mauritiana* | *sechellia* | *melanogaster* | 0.002228 | 0.000000 | 0.002228 | 0.000 | 0.500 | 0.500 | | |
| *ogumai* | *ohnishii* | *montium* | 0.022183 | 0.027833 | -0.005650 | 0.000 | 0.500 | 0.500 | 1 | |
| *paulistorum* | *willistoni* | *willistoni* | 0.018475 | 0.018035 | 0.000440 | 0.250 | 1.000 | 0.750 | | |
| *persimilis* | *pseudoobscura* | *obscura* | 0.013317 | 0.016660 | -0.003343 | 0.000 | 0.500 | 0.500 | | |
| *bogota* | *pseudoobscura* | *obscura* | 0.004841 | 0.016660 | -0.011819 | 0.500 | 0.500 | 0.000 | | |
| *recens* | *subquinaria* | *quinaria* | 0.063736 | 0.066094 | -0.002358 | -0.450 | 0.500 | 0.950 | 2 | *Adhr* not *Adh* |
| *simulans* | *sechellia* | *melanogaster* | 0.027407 | 0.000000 | 0.027407 | 0.000 | 0.500 | 0.500 | | |
| *simulans* | *mauritiana* | *melanogaster* | 0.027407 | 0.002228 | 0.025179 | 0.000 | 0.500 | 0.500 | | |
| *triauraria* | *quadraria* | *montium* | 0.015520 | 0.023228 | -0.007708 | -0.500 | 0.000 | 0.500 | 3, 4 | |

* 12 = species 1 ♀ ✕ species 2 ♂; 21 = species 2 ♀ ✕ species 1 ♂. Data from Table S2 of Yukilevich 2012 Evolution 66: 1430–1446.

**References for estimates of πS:** 1) Chen et al. 2013 Zool. Sci. 30: 1056–1062. 2) Ginsberg et al. 2019 J. Evo. Biol. 32: 1093–1105.
3) Dai, Lu, Lv, Chen, Cheng & Zhang 2001 GenBank: AF348883.1. 4) Watada & Miyake 2011 GenBank: AB669833.1.
All other estimates of πS are summarized in S1 Table.
